# STAT3 regulates the generation of astroglia in human brain organoids with high mTORC1 activity

**DOI:** 10.1101/2023.12.21.572786

**Authors:** Bipan K. Deb, Thomas L. Li, John D. Blair, Dirk Hockemeyer, Helen S. Bateup

## Abstract

During brain development, neural progenitor cells first produce neurons, then astrocytes and other glial cell types, which provide important trophic support and shape neuronal development and function. Intrinsic genetic programs interact with extracellular signals to control progenitor fate, resulting in temporally segregated periods of neurogenesis and gliogenesis. Animal models have implicated STAT3 as an important driver of astrogenesis; however, the signaling pathways that control glial differentiation during human brain development are less well understood. Prior work demonstrated that constitutive activation of mTORC1 signaling in human brain organoid models resulted in the precocious generation of glial-lineage cells. In this study, we tested whether mTORC1 acts via STAT3 to control astrogenesis in brain organoids. We show that knockdown of STAT3 reduces astrogenesis in wild-type organoids and in organoids with constitutively high mTORC1 signaling caused by deletion of the negative regulator TSC2. However, mTORC1 is not required for cytokine-induced activation of STAT3 and expression of the astrocytic protein GFAP. Together, these results show that mTORC1 acts through STAT3 to control astroglia production in human brain organoid models, but that mTOR signaling is dispensable for STAT3-driven astrogenesis.

**Summary statement:** Deb et al, use human brain organoid models to show a requirement for STAT3 downstream of mTORC1 in regulating astrogliogenesis during early human brain development.

## Introduction

The human brain comprises numerous cell types, which include many sub-types of neurons, glia, and other non-neuronal cells. During development, neural progenitor cells (NPCs) first divide symmetrically to generate more progenitors and then asymmetrically to produce intermediate progenitors and neurons (Lui et al., 2011, Silbereis et al., 2016). Following neurogenesis, progenitor cells transition to producing glial-lineage cells, including astrocytes (Allen and Lyons, 2018). The fate switch of NPCs is controlled by intrinsic genetic programs and extracellular signaling cues including cytokines (Miller and Gauthier, 2007). Understanding the signaling pathways that control the initiation of astrogenesis during human brain development has been challenging due to lack of suitable models that can recapitulate human-specific aspects of neural development. Human brain organoids (hBOs) are 3D models of early human brain development that recapitulate important developmental features, including an extended phase of neurogenesis followed by a period of astroglial differentiation and maturation (Sloan et al., 2017, Pasca, 2018). Thus, hBOs recapitulate the time-dependent switch from neurogenesis to gliogenesis and offer a useful experimental model to understand the programs that control this switch (Lanjewar and Sloan, 2021).

Studies in animal models have shown that the mTOR and JAK-STAT signaling pathways play important roles in astrogenesis (Bonni et al., 1997, Hong and Song, 2014, Nakashima et al., 1999, Cloetta et al., 2013, Onda et al., 2002, Fukuda et al., 2007, He et al., 2005). mTOR is part of a conserved signaling pathway that acts as a central regulator of cell growth and metabolism via the control of anabolic and catabolic pathways (Saxton and Sabatini, 2017). mTOR signaling is essential during development (Murakami et al., 2004) and loss of function of either of the two mTOR complexes (mTORC1 and mTORC2) is embryonic lethal in mice (Guertin et al., 2006). A protein complex formed by the TSC1 and TSC2 proteins serves as a critical negative regulator of mTORC1 and loss of function of either of these proteins results in constitutive activation of mTORC1 signaling (Tee et al., 2002). In terms of astrogenesis, studies in mouse models have shown that suppression of mTORC1 leads to a strong impairment in astrocyte production (Cloetta et al., 2013). Conversely, increased mTORC1 signaling due to homozygous deletion of *TSC1* or *TSC2* enhances astroglial differentiation in 2D neural cultures and hBOs (Blair et al., 2018, Grabole et al., 2016, Costa et al., 2016). Recently, it has been shown that gliogenic factors may converge on mTOR signaling to drive astrogenesis in hBOs (Voss et al., 2023). However, the downstream pathways controlled by mTORC1 that regulate the gliogenic switch have not been defined.

STAT proteins comprise a family of transcription factors that regulate tissue-specific gene expression following cytokine receptor-dependent phosphorylation by Janus kinases (JAKs) (Levy and Darnell, 2002). In the context of cancer and the immune system, significant crosstalk between mTOR and STAT3 signaling has been observed (Saleiro and Platanias, 2015, Yokogami et al., 2000, Zhou et al., 2007, Bezzerri et al., 2016). In NPCs, cytokines including LIF (Leukemia Inhibitory Factor) and CNTF (Ciliary Neurotrophic Factor) induce STAT3 phosphorylation and activation (Bonni et al., 1997, Nakashima et al., 1999). Activated STAT3, in complex with other proteins, directly binds to and activates transcription of astroglial genes including *GFAP* (Bonni et al., 1997). Thus, cytokine-mediated activation of STAT3 results in astroglial differentiation from NPCs (Nakashima et al., 1999, Fukuda et al., 2007, Deverman and Patterson, 2009, Mi and Barres, 1999). In hBOs, a LIF-induced increase in STAT3 signaling starting early in development increases the number of GFAP+ astrocytes later in development (Watanabe et al., 2017). Conversely, conditional deletion of STAT3 reduces astroglial differentiation in mouse NPCs (Cao et al., 2010).

Several studies have shown a correlation between high mTORC1 signaling, increased STAT3 phosphorylation, and enhanced glial differentiation (Blair et al., 2018, Onda et al., 2002, Grabole et al., 2016). However, it is unknown whether STAT3 is required for the mTORC1-dependent enhancement of astrogenesis. Here we find that STAT3 knock-down reduces astroglial differentiation in both wild-type (WT) and *TSC2* knock-out (KO) hBOs, which have constitutively high mTORC1 signaling. We additionally observe that LIF and CNTF increase STAT3 phosphorylation and enhance GFAP expression in hBOs, and that this is unaffected by blocking mTORC1 signaling. Together, our results demonstrate that mTORC1 acts through STAT3 to control astrogenesis in hBOs and provide a framework for how mTORC1 signaling interacts with JAK-STAT to control astrocyte differentiation during early human brain development.

## Results

### STAT3 is required for astroglial differentiation in human brain organoids

JAK-STAT is an evolutionarily conserved signaling pathway that can drive astrogenesis in animal models (Miller and Gauthier, 2007). Therefore, we investigated whether STAT3 is required for astroglial differentiation in hBOs. We differentiated WT WIBR3 human embryonic stem cells (hESCs, NIH registry #0079) into brain organoids using an established protocol that recapitulates the time-dependent switch from neurogenesis to gliogenesis, beginning around day 100 post-differentiation (Fig. 1A) (Yoon et al., 2019, Sloan et al., 2017). To reduce STAT3 levels, we used short hairpin RNAs (shRNA) targeting human STAT3 (shSTAT3) that significantly reduced expression when compared to a non-targeting shRNA control (shControl; Fig 1B,C). shSTAT3 or shControl constructs were delivered to WT hBOs on day 35 (D35) of differentiation via lentiviral transduction as part of a vector that co-expresses GFP. At this time point, hBOs primarily contain undifferentiated NPCs (Pasca et al., 2015, Blair et al., 2018). At D100, astroglial or neuronal fate of the shControl or shSTAT3-expressing cells (GFP+) was compared to the neighboring untransduced (GFP-) cells. We found that while shControl expression had no effect on astrocyte differentiation, a smaller percentage of shSTAT3-expressing cells were positive for the astrocyte lineage marker S100B compared to untransduced cells within the same organoid (Fig. 1D-H). This indicates that suppression of STAT3 expression cell autonomously reduces astroglial differentiation.

**Figure 1:**
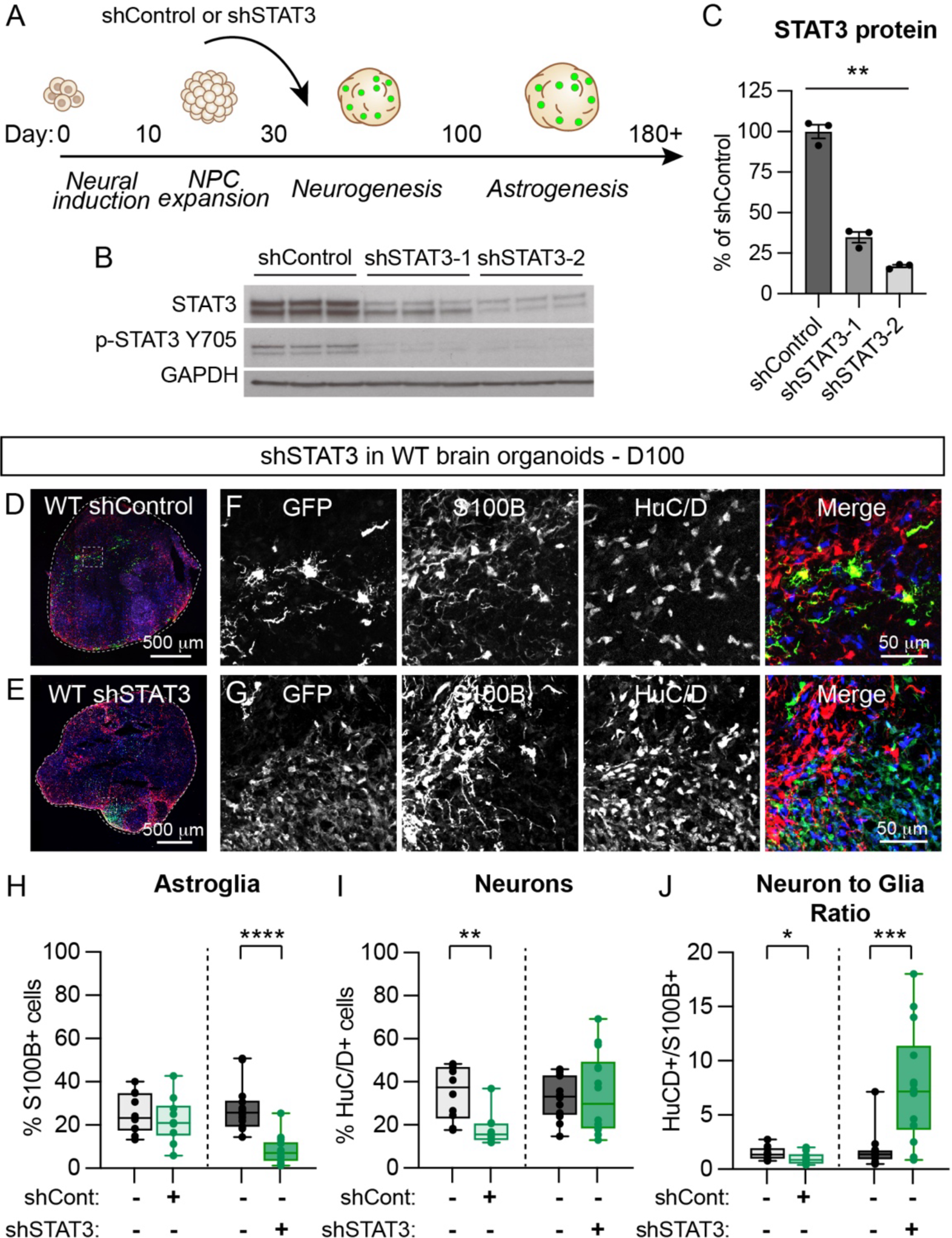
Suppression of STAT3 inhibits astrogenesis in human brain organoids. **(A)** Schematic of the human brain organoid differentiation timeline. Lentiviruses expressing shControl-GFP or shSTAT3-GFP were added on day 35 of differentiation. **(B)** Example western blots of HEK cell lysates treated with shControl or one of two different shSTAT3 lentiviruses. Three independent replicates are shown for each treatment. **(C)** Quantification of total STAT3 protein levels. Bars represent mean +/- SEM, dots represent individual samples. n=3 per condition. **p=0.0036, Kruskal-Wallis test. **(D,E)** Images of day 100 WT whole organoid sections treated with shControl-GFP **(D)** or shSTAT3-GFP **(E)**. **(F)** Zoomed-in images of the boxed region in **D** showing shControl-GFP labeled cells and S100B and HuC/D immunostained cells. **(G)** Zoomed-in images of shSTAT3-GFP cells from **E**. **(H)** Box and whisker plot of the % S100B+ nuclei for organoids from different conditions. shControl (-) vs (+), p=0.6164; shSTAT3 (-) vs (+), ****p<0.0001, Mann-Whitney tests. **(I)** Box and whisker plot of the % HuC/D+ nuclei. shControl (-) vs (+), **p=0.0011; shSTAT3 (-) vs (+), p=0.9820, Mann-Whitney tests. **(J)** Box and whisker plot of the HuC/D+ to S100B+ ratio. shControl (-) vs (+), *p=0.0453; shSTAT3 (-) vs (+), ***p=0.0005, Mann-Whitney tests. For panels **H**, **I**, and **J**, left boxes show GFP- and GFP+ cells from shControl treated organoids. Right boxes show GFP- and GFP+ cells from shSTAT3-2 treated organoids. Boxes extend from the 25^th^ to 75^th^ percentiles, whiskers extend from the min to max values, lines represent the median, and dots represent values for individual organoids. shControl, n=10 organoids; shSTAT3, n=14 organoids.

We examined neuronal differentiation and found that shSTAT3 did not significantly affect the proportion of HuC/D+ neurons generated (Fig. 1I). However, the large reduction in S100B-expressing cells in the shSTAT3 condition significantly increased the neuron-to-glia ratio compared to GFP-cells (Fig. 1J). Notably, we did observe a reduction in the proportion of neurons generated by shControl-expressing cells at D100, compared to neighboring untransduced cells (Fig. 1I), which was unexpected. It is possible that NPCs transduced with lentivirus may show reduced or delayed neuronal differentiation in hBOs. While further work will be needed to assess the effects of STAT3 knock-down on neurogenesis, our data show that reducing STAT3 signaling can suppress astroglial differentiation in hBOs.

### STAT3 knockdown reduces aberrant astroglial differentiation in *TSC2^-/-^* hBOs

Prior studies have reported increased expression of astrocytic proteins, enhanced production of glial-lineage cells, and/or reduced numbers of neurons resulting from constitutive activation of mTORC1 signaling (Blair et al., 2018, Grabole et al., 2016, Onda et al., 2002, Magri et al., 2011, Goto et al., 2011, Costa et al., 2016). We confirmed these results by differentiating WIBR3 hESCs with a CRISPR/Cas9-engineered homozygous deletion of *TSC2* (Blair et al., 2018) into hBOs (Fig. S1A). We verified that *TSC2^-/-^*hBOs have loss of TSC2 protein and constitutive activation of mTORC1 signaling, which can be reversed by treatment with the mTOR inhibitor rapamycin (Fig. S1B-E). We quantified cell type proportions at D100 and found that *TSC2^-/-^*hBOs had an increased proportion of S100B+ glial-lineage cells and a reduced number of HuC/D+ neurons (Fig. S1F-I). This resulted in a strongly reduced neuron-to-glia ratio in *TSC2^-/-^* hBOs (Fig. S1J), consistent with prior reports (Blair et al., 2018).

We had previously shown that homozygous deletion of *TSC1* or *TSC2* increased the levels of phosphorylated STAT3 (pSTAT3-Y705) early during hBO development, prior to the onset of astrogliogenesis (Blair et al., 2018). We therefore hypothesized that STAT3 could act downstream of mTORC1 to drive astroglial differentiation in *TSC2* KO hBOs. To test this, we transduced *TSC2^-/-^* organoids with lentivirus expressing shControl-GFP or shSTAT3-GFP at D35 (Fig. 2A-D). Consistent with our findings in WT hBOs (see Fig. 1), *TSC2*^-/-^ cells expressing shSTAT3 showed a significant reduction in S100B+ astroglia compared to neighboring GFP-cells (Fig. 2E). shSTAT3 did not significantly affect the proportion of HuC/D+ neurons in *TSC2^-/-^* organoids, which remained low across all conditions (Fig. 2F). Similar to WT hBOs, shSTAT3 increased the neuron:glia ratio in *TSC2^-/-^* hBOs, owing to the large reduction in S100B+ cells (Fig. 2G). These effects were replicated using an independent shRNA targeting STAT3 (Fig. S2A-D). Together these data show that STAT3 reduction can effectively reduce the increased astrocyte differentiation due to activation of mTORC1 in hBOs.

**Figure 2:**
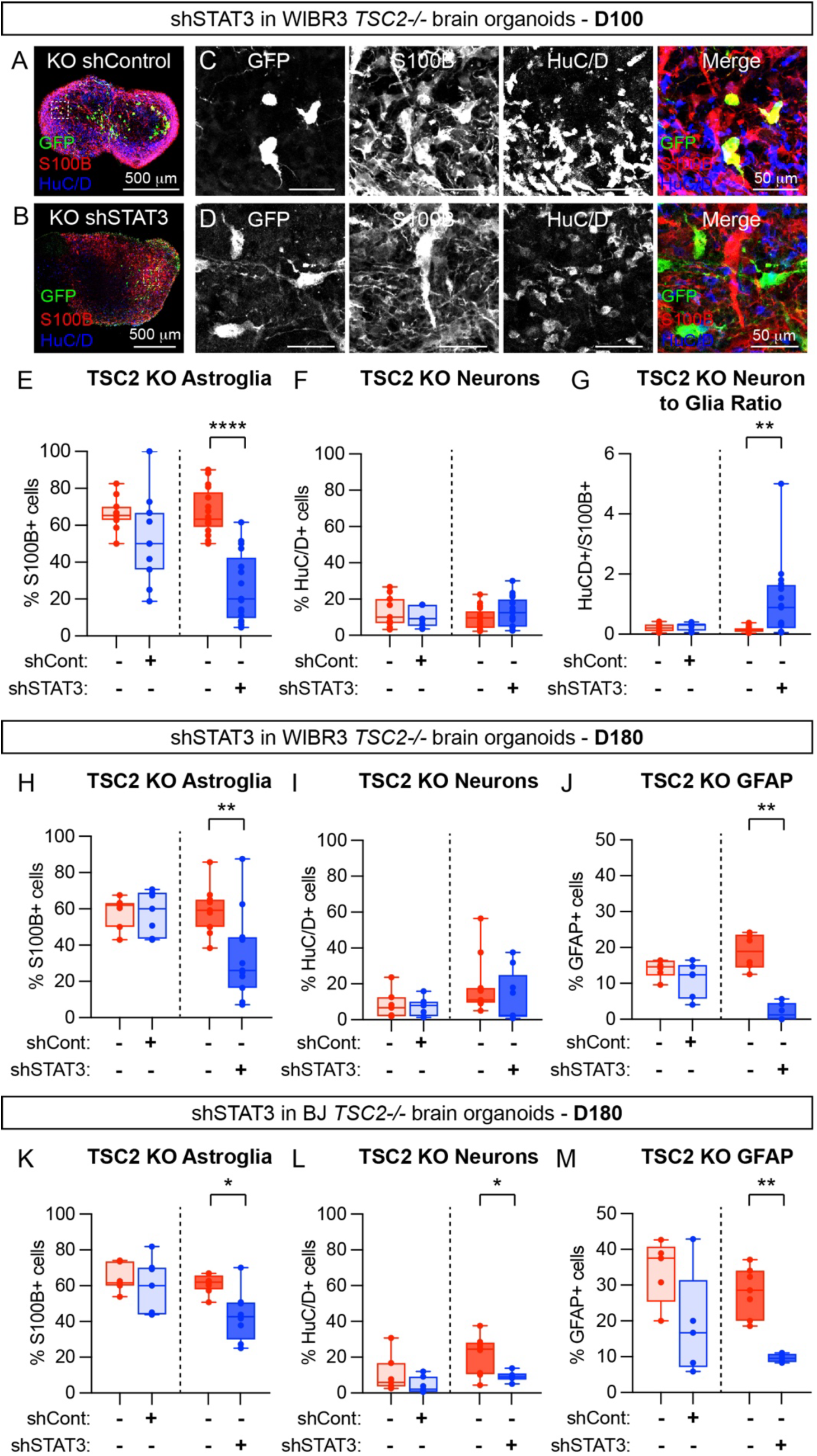
STAT3 reduction prevents the enhanced astrogenesis caused by loss of TSC2. **(A,B)** Images of day 100 *TSC2-/-* whole organoid sections treated with shControl-GFP **(A)** or shSTAT3-GFP **(B)**. **(C)** Zoomed-in images of the boxed region in **A** showing shControl-GFP labeled cells and S100B and HuC/D immunostained cells. **(D)** Zoomed-in images of shSTAT3-GFP cells from **B**. **(E)** Box and whisker plot of the % S100B+ nuclei for D100 *TSC2-/-* WIBR3 organoids from different conditions. shControl (-) vs (+), p=0.1086 (n=11 organoids); shSTAT3 (-) vs (+), ****p<0.0001 (n=17 organoids), unpaired two-tailed t tests. **(F)** Box and whisker plot of the % HuC/D+ nuclei. shControl (-, n=11 organoids) vs (+, n=9 organoids), p=0.4537; shSTAT3 (-) vs (+), p=0.1469 (n=17 organoids), unpaired two-tailed t tests. **(G)** Box and whisker plot of the HuC/D+ to S100B+ ratio. shControl (-, n=11 organoids) vs (+, n=9 organoids), p=0.3384; shSTAT3 (-) vs (+), **p=0.0017 (n=17 organoids), Mann-Whitney tests. For **F** and **G** some organoid sections had zero HuC/D+ shSTAT3-GFP+ cells and were therefore not included in the analysis. **(H)** Box and whisker plot of the % S100B+ nuclei for D180 *TSC2-/-* WIBR3 organoids. shControl (-) vs (+), p=0.9336 (n=7 organoids); shSTAT3 (-) vs (+), **p=0.0049 (n=7 organoids), Mann-Whitney tests. **(I)** Box and whisker plot of the % HuC/D+ nuclei. shControl (-) vs (+), p=0.9318 (n=7 organoids); shSTAT3 (-, n=11 organoids) vs (+, n=9 organoids), p=0.2371, Mann-Whitney tests. **(J)** Box and whisker plot of the % GFAP+ nuclei. shControl (-) vs (+), p=0.3095 (n=6 organoids); shSTAT3 (-) vs (+), **p=0.0022 (n=6 organoids), Mann-Whitney tests. For **I** some organoid sections had zero HuC/D+ shSTAT3-GFP+ cells and were therefore not included in the analysis. **(K)** Box and whisker plot of the % S100B+ nuclei for D180 *TSC2-/-* BJ organoids. shControl (-) vs (+), p=0.4761 (n=7 organoids); shSTAT3 (-) vs (+), *p=0.0134 (n=8 organoids), Mann-Whitney tests. **(L)** Box and whisker plot of the % HuC/D+ nuclei. shControl (-) vs (+), p=0.1282 (n=7 organoids); shSTAT3 (-, n=8 organoids) vs (+, n=6 organoids), *p=0.0426, Mann-Whitney tests. **(M)** Box and whisker plot of the % GFAP+ nuclei. shControl (-) vs (+), p=0.1667 (n=5 organoids); shSTAT3 (-, n=7 organoids) vs (+, n=4 organoids), **p=0.0061, Mann-Whitney tests. For **M** some organoid sections had zero GFAP+ shSTAT3-GFP+ cells and were therefore not included in the analysis. For panels **E-M**, boxes extend from the 25^th^ to 75^th^ percentiles, whiskers extend from the min to max values, lines represent the median, and dots represent values for individual organoids.

### Reduced astroglial differentiation due to STAT3 knockdown is maintained at later stages of development

To investigate if the reduced proportion of S100B+ cells observed after STAT3 knockdown signified a delay in astrocyte differentiation or a sustained reduction, we maintained *TSC2*^-/-^ hBOs transduced with either shControl-GFP or shSTAT3-GFP for 6 months and investigated cell fate at day 180 post-differentiation (D180). Similar to D100, we found that shSTAT3-transduced cells had a significant reduction in the proportion of S100B+ cells at D180 (Fig. 2H and Fig. S2E,F), with no effect on the proportion of HuC/D+ cells (Fig. 2I). To confirm that this reflected suppression of astroglial fate and not a reduction of S100B expression, we examined another canonical marker of astrocytes, GFAP, which is expressed in a sub-set of more mature cells present at this stage of differentiation (Fig. S2G,H). We found a strong reduction in the proportion of GFAP+ astroglia differentiated from shSTAT3-transduced *TSC2^-/-^* cells (Fig. 2J), demonstrating that shSTAT3 suppresses multiple markers of astroglial differentiation at later stages of hBO development.

To confirm these findings, we tested the effects of STAT3 knockdown in *TSC2*^-/-^ hBOs derived from an independent hPSC line. We used an hiPSC line reprogrammed from BJ fibroblasts, genetically engineered to have homozygous deletion of *TSC2* (Blair et al., 2018). shSTAT3 significantly reduced the percentage of S100B+ cells at D180 in BJ *TSC2*^-/-^ organoids (Fig. 2K), with a small effect on the proportion of HuC/D+ cells (Fig. 2L and Fig. S2I,J). The proportion of GFAP+ cells was also strongly decreased by shSTAT3 in D180 BJ *TSC2^-/-^*organoids (Fig. 2M and Fig. S2K,L). Together, these data show that STAT3 reduction in *TSC2*^-/-^ hBOs causes a sustained inhibition of astroglial differentiation.

### STAT3 knockdown does not reverse mTORC1 hyperactivity

Our results indicated that suppression of STAT3 could counteract the pro-gliogenic effects of high mTORC1 signaling in hBOs. To determine whether this was due to suppression of mTORC1 signaling by shSTAT3, we measured two canonical read-outs of mTORC1 signaling, the phosphorylation of ribosomal protein S6 (p-RPS6) and cell size. We found that shSTAT3 did not significantly affect p-RPS6 levels in either WIBR3 WT or *TSC2^-/-^* hBOs at D100, although p-RPS6 levels were elevated in *TSC2* KO cells, as expected (Fig. S3A-E). Additionally, shSTAT3 did not reverse the cellular hypertrophy of HuC/D+ neurons or S100B+ astroglia in *TSC2^-/-^* hBOs (Fig. S3F,G). These data indicate that STAT3 acts downstream of mTORC1 to control glial differentiation and that canonical consequences of mTORC1 hyperactivation are not strongly affected by STAT3 suppression.

### Cells with STAT3 knockdown maintain expression of SOX2

We next investigated whether the failure of STAT3 knock-down cells to commit to an astroglial fate reflected prolonged maintenance of a progenitor state. This possibility was supported by the observation that many shSTAT3 transduced cells expressed neither S100B nor HuC/D, which was the case for both WT and *TSC2^-/-^* organoids (Fig. 3A-D). Although this was also observed in some shControl-transduced cells (Fig. 3B), the effect was transient and was no longer apparent by day 180 (Fig. 3C). To test whether the shSTAT3-transduced cells could be progenitors, we immunostained D100 WT and *TSC2^-/-^* hBOs for SOX2, a transcription factor expressed in NPCs but not in differentiated neurons (Pevny and Placzek, 2005) (Fig. 3E,F). We found that significantly more shSTAT3-expressing cells were positive for SOX2 in both WT and *TSC2^-/-^* organoids (Fig. 3G,H). These data suggest that at day 100, a large proportion of shSTAT3-expressing cells were likely progenitors.

**Figure 3:**
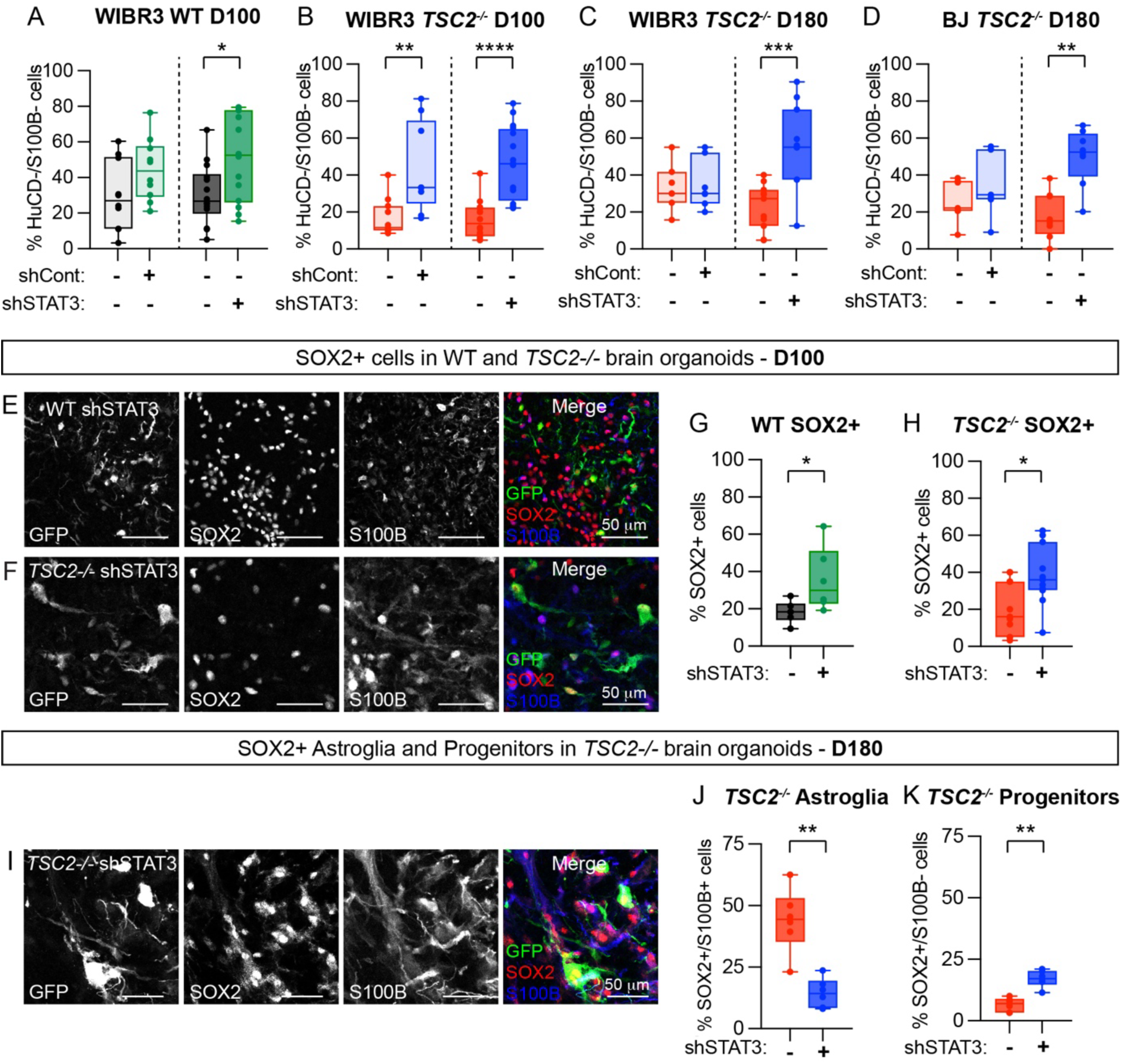
STAT3 reduction leads to a maintenance of uncommitted progenitor cells. **(A-D)** Box and whisker plots of the % HuC/D-/S100B-nuclei per organoid from different conditions. Left boxes show GFP- and GFP+ cells from shControl treated organoids. Right boxes show GFP- and GFP+ cells from shSTAT3-2 treated organoids. Boxes extend from the 25^th^ to 75^th^ percentiles, whiskers extend from the min to max values, lines represent the median, and dots represent values for individual organoids. **(A)** WIBR3 WT day 100, shControl (-) vs (+), p=0.1230 (n=10 organoids); shSTAT3 (-) vs (+), *p=0.0241 (n=14 organoids), Mann-Whitney tests. **(B)** WIBR3 *TSC2-/-* day 100, shControl (-, n=11 organoids) vs (+, n=9 organoids), **p=0.0031; shSTAT3 (-) vs (+), ****p<0.0001 (n=16 organoids), Mann-Whitney tests. **(C)** WIBR3 *TSC2-/-* day 180, shControl (-) vs (+), p=0.9353 (n=7 organoids); shSTAT3 (-) vs (+), ***p=0.0004 (n=11 organoids), Mann-Whitney tests. **(D)** BJ *TSC2-/-* day 180, shControl (-) vs (+), p=0.2593 (n=7 organoids); shSTAT3 (-) vs (+), **p=0.0019 (n=8 organoids), Mann-Whitney tests. **(E)** Example images of WT shSTAT3-GFP expressing cells in D100 organoid sections immunostained for SOX2 and S100B. **(F)** Example images of *TSC2-/-* shSTAT3-GFP expressing cells in D100 organoid sections. **(G)** Box and whisker plot of the % SOX2+ nuclei in WT D100 organoids treated with shSTAT3. *p=0.0260 (n=6 organoids), Mann-Whitney test. **(H)** Box and whisker plot of the % SOX2+ nuclei in *TSC2-/-* D100 organoids treated with shSTAT3. *p=0.0137 (n=11 organoids), Mann-Whitney test. **(I)** Example images of *TSC2-/-* shSTAT3-GFP expressing cells in D180 organoid sections immunostained for SOX2 and S100B. **(J)** Box and whisker plot of the % SOX2+/S100B+ nuclei in *TSC2-/-* D180 organoids treated with shSTAT3; **p=0.0043 (n=6 organoids per condition), Mann-Whitney test. **(K)** Box and whisker plot of the % SOX2+/S100B-nuclei in *TSC2-/-* D180 organoids; **p=0.0022 (n=6 organoids), Mann-Whitney test.

We next investigated if the shSTAT3-transduced cells remained as progenitors at later stages of development. Since SOX2 continues to be expressed in differentiated astrocytes (Wang et al., 2022), we co-stained hBOs with SOX2 and S100B to distinguish undifferentiated progenitors (SOX2+/S100B-) from cells committed to an astroglial fate (SOX2+/S100B+) (Fig. 3I). Consistent with reduced astroglial differentiation, fewer shSTAT3-transduced *TSC2^-/-^* cells were SOX2+/S100B+ compared to neighboring untransduced cells (Fig. 3J). By contrast, shSTAT3 led to a doubling of the proportion of SOX2+/S100B-*TSC2^-/-^* cells (Fig. 3K), supporting the idea that STAT3 suppression leads to a maintained progenitor state. From this data we conclude that STAT3 knockdown results in prolonged maintenance of neural progenitors, which fail to acquire an astroglial fate by 6 months in hBOs.

### STAT3 activation induces GFAP independent of mTORC1

Our results indicate that increasing mTORC1 signaling boosts astroglial production while suppressing STAT3 signaling decreases it. To determine if these effects are bidirectional, we treated WT hBOs from day 35 onwards with the mTOR inhibitor rapamycin or the cytokines LIF and CNTF to activate STAT3 (Bonni et al., 1997) (Fig. 4A). We analyzed hBOs at D120 and observed that rapamycin reduced overall organoid size, while LIF/CNTF tended to increase it, compared to controls (Fig. 4B,C). When rapamycin was applied together with LIF/CNTF, hBO size was smaller than controls, indicating that mTOR suppression had a dominant effect on organoid size (Fig. 4B,C). We assessed the levels of p-RPS6 by western blot and found the expected reduction by rapamycin, with no significant change due to LIF/CNTF treatment, again demonstrating that modulation of STAT3 activation does not strongly affect mTORC1 signaling (Fig. 4D,E).

**Figure 4:**
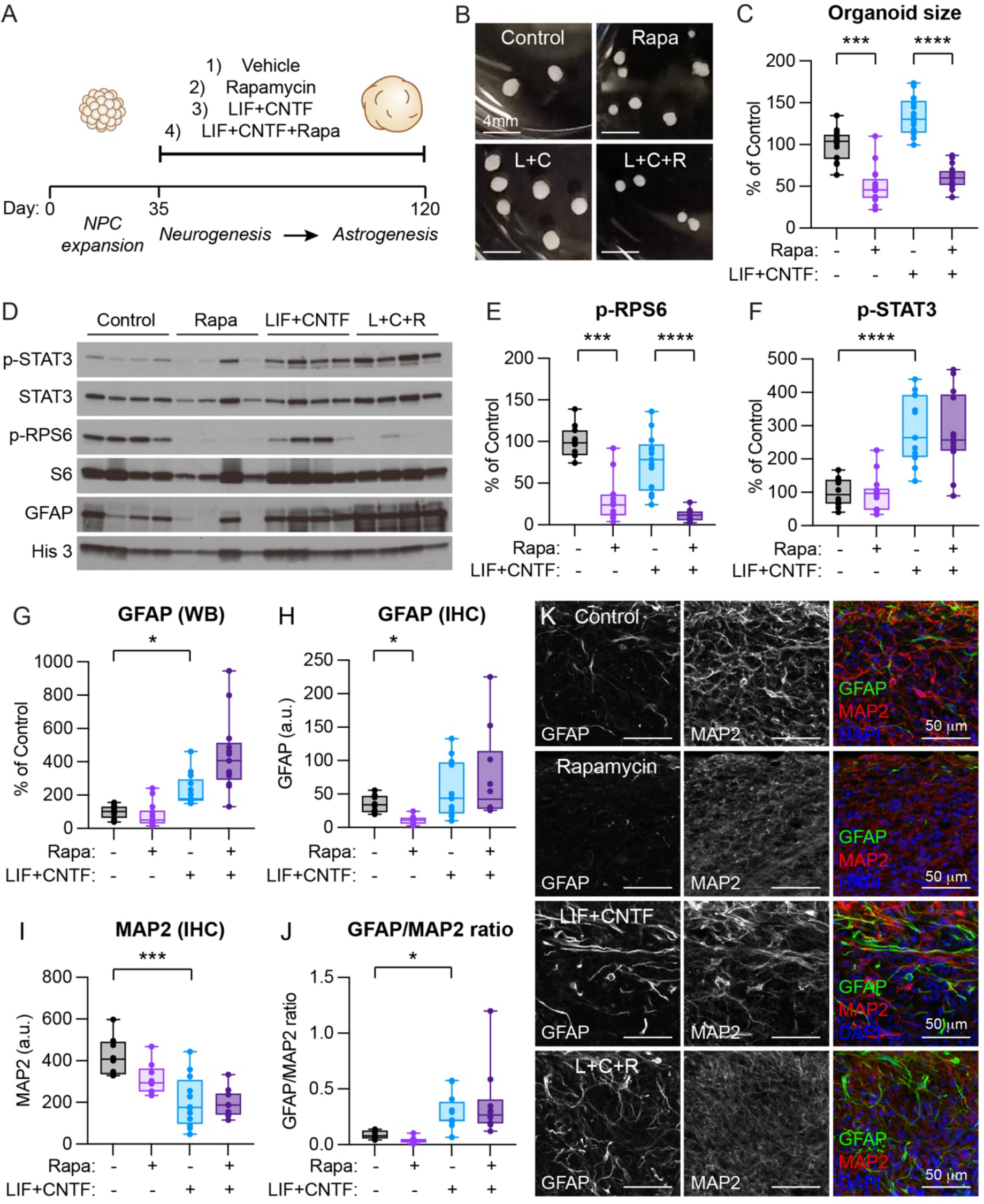
mTOR signaling is not required for p-STAT3 and GFAP induction by LIF/CNTF. **(A)** Schematic of the experiment. **(B)** Example images of whole organoids in culture from the indicated groups. L+C = LIF+CNTF, L+C+R = LIF+CNTF+Rapamycin. All scale bars = 4 mm. **(C)** Box and whisker plot of organoid area from the indicated treatment groups expressed as a percent of control. ****p<0.0001, Kruskal-Wallis test; Control (n=14 organoids) vs Rapa (n=17 organoids), ***p=0.0002, Control vs LIF+CNTF (n=17 organoids) p=0.1163, LIF+CNTF vs LIF+CNTF+Rapa (n=17 organoids) ****p<0.0001, Dunn’s multiple comparisons test. **(D)** Representative western blots for the antibodies shown. Four independent samples per treatment are shown. **(E)** Box and whisker plot of p-RPS6 (Ser240/244) measured by western blot, normalized to total RPS6. ****p<0.0001, Kruskal-Wallis test; Control (n=11 organoids) vs Rapa (n=14 organoids), ***p=0.0004, Control vs LIF+CNTF (n=13 organoids) p=0.7798, LIF+CNTF vs LIF+CNTF+Rapa (n=13 organoids) ****p<0.0001, Dunn’s multiple comparisons test. **(F)** Box and whisker plot of p-STAT3 (Y705) measured by western blot, normalized to total STAT3. ****p<0.0001, Brown-Forsythe ANOVA test; Control (n=12 organoids) vs Rapa (n=14 organoids), p=0.9943, Control vs LIF+CNTF (n=13 organoids) ****p<0.0001, LIF+CNTF vs LIF+CNTF+Rapa (n=13 organoids) p=0.9976, Dunnett’s T3 comparisons test. **(G)** Box and whisker plot of GFAP measured by western blot, normalized to Histone 3. ****p<0.0001, Kruskal-Wallis test; Control vs Rapa, p>0.9999, Control vs LIF+CNTF *p=0.0136, LIF+CNTF vs LIF+CNTF+Rapa p=0.3343, Dunn’s multiple comparisons test (n is the same as for panel **F**). **(H)** Box and whisker plot of the thresholded GFAP signal per nuclei measured by IHC. ***p=0.0004, Kruskal-Wallis test; Control (n=8 organoids) vs Rapa (n=9 organoids), *p=0.0143, Control vs LIF+CNTF (n=12 organoids) p>0.9999, LIF+CNTF vs LIF+CNTF+Rapa (n=10 organoids) p>0.9999, Dunn’s multiple comparisons test. **(I)** Box and whisker plot of the thresholded MAP2 signal per nuclei measured by IHC. ***p=0.0003, Kruskal-Wallis test; Control vs Rapa, p=0.3532, Control vs LIF+CNTF ***p=0.0007, LIF+CNTF vs LIF+CNTF+Rapa p>0.9999, Dunn’s multiple comparisons test (n is the same as for panel **H**). **(J)** Box and whisker plot of the GFAP to MAP2 ratio per organoid measured by IHC. ****p<0.0001, Kruskal-Wallis test; Control vs Rapa, p=0.5695, Control vs LIF+CNTF *p=0.0172, LIF+CNTF vs LIF+CNTF+Rapa p>0.9999, Dunn’s multiple comparisons test (n is the same as for panel **H**). **(K)** Example images of organoid sections from the indicated treatment groups showing GFAP and MAP2 immunostaining. Nuclei are labeled with DAPI in blue in the merged images. For panels C and E-J, boxes extend from the 25^th^ to 75^th^ percentiles, whiskers extend from the min to max values, lines represent the median, and dots represent values for individual organoids.

We examined p-STAT3 levels and found that LIF/CNTF significantly increased the ratio of p-STAT3 to total STAT3, as expected (Fig. 4F). Notably, co-administration of rapamycin with LIF/CNTF did not affect the induction of p-STAT3 (Fig. 4F), signifying that mTOR is not required for cytokine-mediated STAT3 activation. We measured GFAP levels in these organoids and found that LIF/CNTF significantly increased GFAP protein expression (Fig. 4G). Consistent with the p-STAT3 results, rapamycin did not block the induction of GFAP by LIF/CNTF, and, in some cases, potentiated it (Fig. 4G).

We further investigated these effects in a separate batch of organoids immunostained for GFAP and the neuronal dendritic protein MAP2. In this experiment, we found that rapamycin reduced GFAP expression, consistent with a progliogenic effect of mTOR signaling (Fig. 4H,K). Interestingly LIF+CNTF treatment significantly reduced MAP2 levels suggesting reduced neuronal differentiation (Fig. 4I,K). We calculated the ratio of GFAP to MAP2 for each organoid, and found a significant increase with LIF+CNTF treatment, which was not affected by rapamycin (Fig. 4J,K). Together, these results show that mTOR signaling is not required for p-STAT3 and GFAP induction by cytokines.

### Conclusions

In this study we show that mTORC1 and JAK-STAT are important regulators of astrogenesis in hBO models. We demonstrate that STAT3 signaling is required for normal astroglial differentiation in hBOs. In addition, we show that aberrant gliogenesis induced by constitutively active mTORC1 can be reversed by suppression of STAT3 expression. These findings have implications for neurodevelopmental disorders, specifically mTORopathies, which are caused by mutations that lead to overactive mTORC1 signaling and result in brain malformations (Blair and Bateup, 2020, Karalis and Bateup, 2021). However, mTOR activity appears to be dispensable for mediating astrogenesis downstream of cytokine-induced STAT3 signaling. Open areas for future studies include determining the relative contribution of mTORC1 signaling to gliogenesis under physiological conditions, the mechanism(s) by which mTORC1 signaling induces STAT3 phosphorylation, and how manipulation of STAT3 activation impacts neurogenesis in hBOs. These remaining questions are summarized together with our main findings in Fig. S4.

## Materials and Methods

### hESC and hiPSC cultures

Human stem cell use was approved by the University of California, Berkeley Stem Cell Research Oversight committee. WT WIBR3 (NIH registry #0079) hESCs and BJ hiPSCs (Blair et al., 2018) were used in this study. *TSC2*^-/-^ hESCs and hiPSCs were generated previously by CRISPR/Cas9 editing using two sgRNAs flanking exon 5 of *TSC2* (Blair et al., 2018). Homozygous deletion of exon 5 results in complete loss of TSC2 protein expression (Blair et al., 2018).

All hESC and hiPSC lines were maintained on a layer of inactivated mouse embryonic fibroblasts (MEFs) in hPSC medium composed of DMEM/F12 (Thermo Fisher: 11320-033) supplemented with 20% KnockOut Serum Replacement (KSR) (Thermo Fisher: 10828028), 2 mM glutamine (Thermo Fisher: 25030-081), 1% non-essential amino acids (Thermo Fisher: 11140-050), 0.1 mM β-mercaptoethanol (Thermo Fisher: 21985-023), 1000 U/ml penicillin/streptomycin (Thermo Fisher: 15140-122), and 4 ng/ml FGF2 (Thermo Fisher: PHG0261). Cultures were passaged every 7 days with collagenase type IV (Thermo Fisher: 17104019, 1.5 mg/ml) and gravitational sedimentation by washing 3 times in wash media composed of DMEM/F12 supplemented with 5% fetal bovine serum (Thermo Fisher: 10082– 147) and 1000 U/ml penicillin/streptomycin. All hPSC lines were tested monthly for mycoplasma contamination.

### Array Comparative Genomic Hybridization (aCGH) analysis

Genomic integrity of the stem cell lines was confirmed using array comparative genomic hybridization analysis (Table S1). For sample preparation, collagenase (1.5 mg/ml) was used to separate the hPSC colonies from MEF feeders. The colonies were subsequently centrifuged at 1000 rpm for 5 mins. The cell pellet obtained was frozen and sent to Cell Line Genetics (Madison, WI) for analysis. For aCGH, >1 μg of DNA was extracted from the samples and run on an Agilent SurePrint G3 Human CGH Microarray covering 60,000 probes evenly spaced across the genome.

### Human brain organoid generation

Differentiation of hESCs and hiPSCs into three-dimensional hBOs was performed according to a modified version of an established protocol (Yoon et al., 2019), using stem cells cultured on a MEF feeder layer (Blair et al., 2018). Undifferentiated colonies of hESCs or hiPSCs were removed from the MEF feeder layer using collagenase (1.5mg/mL). Stem cell colonies were washed once with hPSC media and suspended in hPSC media without FGF2 (hPSC-FGF2 media), supplemented with 10 μM Y-27632 (Chemdea: CD0141) and plated into a single well of a 6-well Aggrewell plate (Stemcell Technologies Aggrewell 800) that was coated with 2 ml of anti-adherence rinsing solution (Stem cell technologies: 07010) for 5 mins before plating the cells. Overnight, aggregates were transferred from the Aggrewell to an ultra-low attachment (ULA) 6-well plate (Corning 3471). On days 1-5, the embryoid bodies were transferred to hPSC media without FGF2, supplemented with 10 μM Dorsomorphin (Abcam: ab146597) and 10 μM SB431542 (Fisher scientific: 501360888) in 6-well low attachment plates (Corning: 3471). On day 6, the developing hBOs were transferred to neural induction media composed of Neurobasal-A (Thermo Fisher: 10888022), B-27 Supplement without Vitamin A (Thermo Fisher: 12587-010), PenStrep (Thermo Fisher: 15140-122), and Glutamax (Life Technologies: 35050061), supplemented with 20 ng/ml FGF (Life Technologies corporation: PHG0261) and 20 ng/ml EGF (R and D systems: 236EG200). Media was subsequently changed every day from days 6-15 and then every other day until day 25. From days 25-43, the developing hBOs were grown in neural induction media supplemented with 20 ng/ml BDNF (VWR: RL009-001-C27) and 20 ng/ml NT-3 (Sigma: SRP3128-10UG), and media was changed every 4 days. From day 43 onwards, hBOs were maintained in neural induction media without BDNF or NT-3, with media changes every 4 days until harvested. At this stage, each hBO was transferred to a single well of a 24-well ULA plate (Corning: 3473) and maintained this way for the remainder of the differentiation.

### shRNA and lentivirus production

shRNA sequences targeting human *STAT3* were obtained from the Broad Institute GPP Web Portal (https://portals.broadinstitute.org/gpp/public/gene/search). The sequences of the hairpin oligonucleotides used for generating each shRNA construct are as follows:

shSTAT3-2(TRCN0000020842):

Forward: 5’-CCGG**GCACAATCTACGAAGAATCAA**CTCGAGTTGATTCTTCGTAGATTGTGCTTTTTG-3’,

Reverse: 5’-AATTCAAAAA**GCACAATCTACGAAGAATCAA**CTCGAGTTGATTCTTCGTAGATTGTGC-3’

shSTAT3-1(TRCN0000071455):

Forward: 5’-CCGG**CACCATTCATTGATGCAGTTT**CTCGAGAAACTGCATCAATGAATGGTGTTTTTG-3’,

Reverse: 5’-AATTCAAAAA**CACCATTCATTGATGCAGTTT**CTCGAGAAACTGCATCAATGAATGGTG-3’

The 21bp sequences targeted by the shRNA constructs are highlighted in bold. The 5’ end of each oligonucleotide contains the overhang for Age1 and the 3’ end contains the overhang for EcoR1.

To generate lentiviral expression constructs, we used pKLO.1 containing a non-targeting sequence (Addgene #1864) (Sarbassov et al., 2005) and cloned in GFP in place of the puromycin resistance cassette (shControl-GFP). For the shSTAT3 constructs, the forward and reverse oligonucleotides were annealed and inserted into the pKLO.1-GFP vector using EcoR1 and Age1 (shSTAT3-GFP).

For the generation of lentiviral particles, the above plasmids were transfected together with lentiviral packaging plasmids expressing VsVg, RRE and REV (Addgene #12253, 12251, and 12259) into HEK cells. GFP expression could be detected 24 hours after transfection. To collect the lentiviral particles, the culture medium of the transfected cells was collected 72h post-transfection and mixed with LentiX Concentrator (Takara Biosciences: 631232) according to the manufacturer’s protocol. The mixture was spun down after an overnight incubation at 4 degrees C and the viral pellet was resuspended in DMEM/F-12 medium. The resuspended virus was stored at -80 degrees C until further use. Viruses were generated twice (two batches) using the same protocol.

### Lentiviral transduction

For lentiviral transduction on day 35 of differentiation, 4-6 hBOs were put into a single well of a 6-well ULA plate in neural induction media comprising Neurobasal (-A) media supplemented with B27 (-A), PenStrep and Glutamax. 5 μl of either the shControl, shSTAT3-1, or shSTAT3-2 lentivirus was added to the media. Media was changed the next day and hBOs were maintained as described earlier. Viral transduction was sub-saturating to allow for transduced GFP+ and untransduced GFP-cells to be compared within the same organoid.

### Western blotting

hBOs were harvested in lysis buffer containing 1% SDS, Halt phosphatase inhibitor (Thermo Fisher: PI78420), and protease inhibitor (Roche: 4693159001) in 1× PBS. Total protein was determined by a BCA assay (Thermo Fisher: PI23227) and 10 μg of protein in Laemmli sample buffer (Bio-Rad:161-0747) were loaded onto 4-15% Criterion TGX gels (Bio-Rad: 5671084). Proteins were transferred to PVDF membranes (Bio-Rad: 1620177), blocked in 5% milk in TBS-Tween for one hour at room temperature (RT), and incubated with primary antibodies diluted in 5% milk in TBS-Tween overnight at 4°C. The next day, membranes were incubated with HRP-conjugated secondary antibodies (1:5000, Bio-Rad: Immune-star goat anti-rabbit HRP conjugate: 170-5046 or Immune-star goat anti-mouse HRP conjugate: 1705047) for one hour at RT, washed in TBS-Tween, incubated with chemiluminesence substrate (Perkin-Elmer: NEL105001EA), and then developed on GE Amersham Hyperfilm ECL (VWR: 95017-661). Membranes were stripped with 6M guanidine hydrochloride (Sigma: G3272) to re-blot on the following days.

Band intensity was quantified by densitometry using Image J software. Phospho-proteins were normalized to their respective total proteins. Non-phospho-proteins were normalized to Histone 3, which was used as the loading control. Primary antibodies used were in the following dilutions: Rabbit anti-phospho-STAT3-Y705 (CST: 9145, 1:500), Mouse anti-STAT3 (CST: 9139, 1:1000), Rabbit anti-phospho-RPS6-S240/44 (CST: 5364S, 1:1000), Rabbit anti-RPS6 (CST: 2317S, 1:1000), Rabbit anti-phospho-AKT-S473 (CST:4060, 1:1000), Rabbit anti-AKT (CST: 4691, 1:1000), Rabbit anti-GFAP (Thermo Fisher: PIPA585109, 1:2000), Mouse anti-Histone 3 (CST:3638S, 1:2500).

### Immunohistochemistry (IHC)

Organoids were removed from the culture media, washed once in ice-cold 1× PBS with Ca^2+^/Mg^2+^ and then fixed in 4% paraformaldehyde (PFA, VWR: 100504-940) for 30 mins at room temperature. After fixation, hBOs were transferred to a conical tube containing 30% sucrose solution in 1X PBS and allowed to settle overnight at 4°C. The following day, hBOs were frozen in tissue blocks with OCT compound (Fisher: 23-730-571) and sectioned on a cryostat (Microm HM550) into 16 μm sections. hBO sections were washed once with 1x PBS and blocked in blocking buffer containing 10% normal goat serum (NGS, Vector labs: S-1000-20), 0.1% BSA and 0.3% Triton-X in 1× PBS for one hour at RT. hBO sections were subsequently incubated overnight at 4°C in primary antibodies in antibody dilution buffer (2% NGS and 0.1% Triton-X in 1× PBS). The next day, hBO sections were washed three times with 1× PBS and incubated in secondary antibodies (1:500) in antibody dilution buffer for one hour at RT. For all experiments, Hoechst (Thermo Fisher: 62249) or DAPI was added along with the secondary antibody solution to stain nuclei in the organoid sections. hBO sections were washed again three times with 1× PBS. Slides were mounted with ProLong Gold Antifade mountant without DAPI (Thermo Fischer: 36935) and allowed to set for one day before imaging. Primary antibodies were used in the following dilutions: Mouse anti-HuC/D (Fisher: A21271, 1:500), Rabbit anti-S100B (Abcam: ab868, 1:300), Chicken anti-GFP (AbCam: ab13970, 1:1000), Rabbit anti-GFAP (Thermo Scientific: PIPA585109, 1:300), Mouse anti-SOX2 (AbCam: ab97959-100ug, 1:250) and Chicken anti-MAP2 (Abcam ab5392, 1:5000). The following secondary antibodies were used at 1:500 dilution: Alexa Fluor donkey anti-chicken 488 (Thermo Fisher: A78948), Alexa Fluor donkey anti-rabbit 546 (Thermo Fisher: A-10040) and Alexa Fluor goat anti-mouse 633 (Thermo Fisher: A-21050).

### Confocal microscopy and image analysis

Images were acquired on a FV3000 Olympus Fluoview confocal microscope with 10× (Olympus: UPLXAPO10X) or 20× objectives (Olympus: UCPLFLN20X) using Fluoview 3000 software. For experiments in which multiple conditions were quantitatively compared, the same acquisition settings were used for each image. All confocal images were processed using Image J. Only healthy regions of the spheroids, defined by intact Hoechst+ nuclei, were used for quantification and analysis. To quantify the percentage of shSTAT3 or shControl transduced GFP+ cells that were positive for a neuronal, glial or progenitor marker, ROIs were drawn manually around the GFP+ cells in Image J. For quantifying the presence or absence of a cellular marker, a threshold was set based on the background fluorescence level in Image J and applied to all ROIs. If the mean fluorescence units averaged across the ROI were above this threshold, the cell was considered positive for the marker. GFP-cells identified using intact Hoechst staining were selected in the immediate vicinity of the GFP+ cells. ROIs were drawn around these GFP-nuclei and presence or absence of a cellular marker was quantified similarly as described above for the GFP+ cells. To quantify the cell body size, ROIs were manually drawn around each cell in Image J. ROIs were saved and applied to the p-RPS6 images where mean fluorescence units averaged across the ROI were used as the p-RPS6 value for that cell. The number of hBOs used for quantification and the number of cells counted are indicated in the figure legends.

### Cytokine and rapamycin treatment

20 ug/ml stock solutions of Leukemia inhibitory factor (LIF, EMD Millipore: GF342) and Ciliary Neurotrophic factor (CNTF, Gibco: PHC7015) were prepared in ultrapure sterile water and added to a final concentration of 20 ng/ml each to the organoid culture media comprising Neurobasal (-A) media supplemented with B27 (-A), PenStrep and Glutamax. Rapamycin (LC Laboratories: R-5000) was diluted in EtOH and added to a final concentration of 50 nM to the organoid culture media. Four different treatment combinations were used: Vehicle (EtOH), Rapamycin, LIF+ CNTF, LIF+ CNTF + Rapamycin. All treatments were started at day 35 and subsequently added with each media change for the remainder of the differentiation until day 120 when the hBOs were harvested for either Western blot or cryopreserved for IHC.

### Organoid size quantification

For quantifying organoid size, organoids were removed from their growth medium and washed with 1X PBS. All organoids of a particular treatment condition were placed in individual wells of an ultra-low attachment 6 well plate in 1X PBS and an image of all wells containing organoids of all treatment conditions in a single field of view was obtained using a Motorola Moto G power (2022) camera. This image was opened using ImageJ and an ROI was drawn along the outer boundary of each individual organoid to compute the area.

### GFAP and MAP2 IHC quantification

Organoids were sectioned and immunostained for GFAP and MAP2 as described above. Whole organoid sections were imaged with an Olympus FV3000 confocal microscope using a 20x objective. Quantifications were performed using the Stardist (Weigert et al., 2020) and Scikit-Image (van der Walt et al., 2014) python packages. For each organoid, all nuclei were segmented using Stardist to generate a nuclei label image. Debris was filtered using a size threshold, and labels corresponding to nuclei in adjacent organoids were manually removed. The MAP2 and GFAP channels were binarized using Otsu’s method (skimage.filters.threshold_otsu) across the full organoid. Label dilation and erosion manipulations were performed on the nuclei label image to generate a mask that included the outermost 155 µm (500 px) region of each organoid. To calculate the MAP2 and GFAP values for each organoid, the total number of above-threshold pixels within the masked region was summed, and then divided by the total number of nuclei within the masked region. This value corresponds to an average MAP2+ or GFAP+ area per nucleus.

### Statistics summary

Sample size was determined based on previous studies. All samples that passed technical quality control were included in the statistical analysis. The proportion of cells that were positive for a cellular marker was compared between GFP+ and GFP-cells within the same organoid to account for any variability in differentiation. Organoids were randomly assigned to different treatment conditions. Quantifications were not performed blindly. Statistical analysis was performed using GraphPad Prism software (version 10) and the specific test for each experiment is noted in the figure legend. The means of two normally distributed groups were compared using an unpaired two-tailed t-test. To compare the means of three or more normally distributed groups, a one-way ANOVA was used followed by Sidak’s multiple comparisons test. For data sets with non-normal distributions, a Mann-Whitney test or Kruskal-Wallis test was used followed by Dunn’s multiple comparisons test. Exact P values are reported in the figure legends. P values were corrected for multiple comparisons.

## Acknowledgements

We thank members of the Bateup and Hockemeyer labs for technical training, technical support, and feedback regarding this work.

## Competing interests

No competing interests declared.

## Funding

This work was supported by a National Institutes of Health grant (#R01NS097823) to H.S.B.. B.K.D. was supported by a TS Alliance Postdoctoral Fellowship Grant Award (#01-20) and a postdoctoral fellowship from the Siebel Stem Cell Institute. T.L.L. is a CIRM Scholar of the CIRM Training Program EDUC4-12790. J.D.B. was supported by a Frederick Banting and Charles Best Canada Graduate Scholarship from Canadian Institutes for Health Research (#356733). H.S.B. and D.H. are Chan Zuckerberg Biohub Investigators. H.S.B. is a Weill Neurohub Investigator.

## Data availability statement

All relevant data can be found within the article and its supplementary information.

## Supplementary Information

**Supplementary Figure 1:**
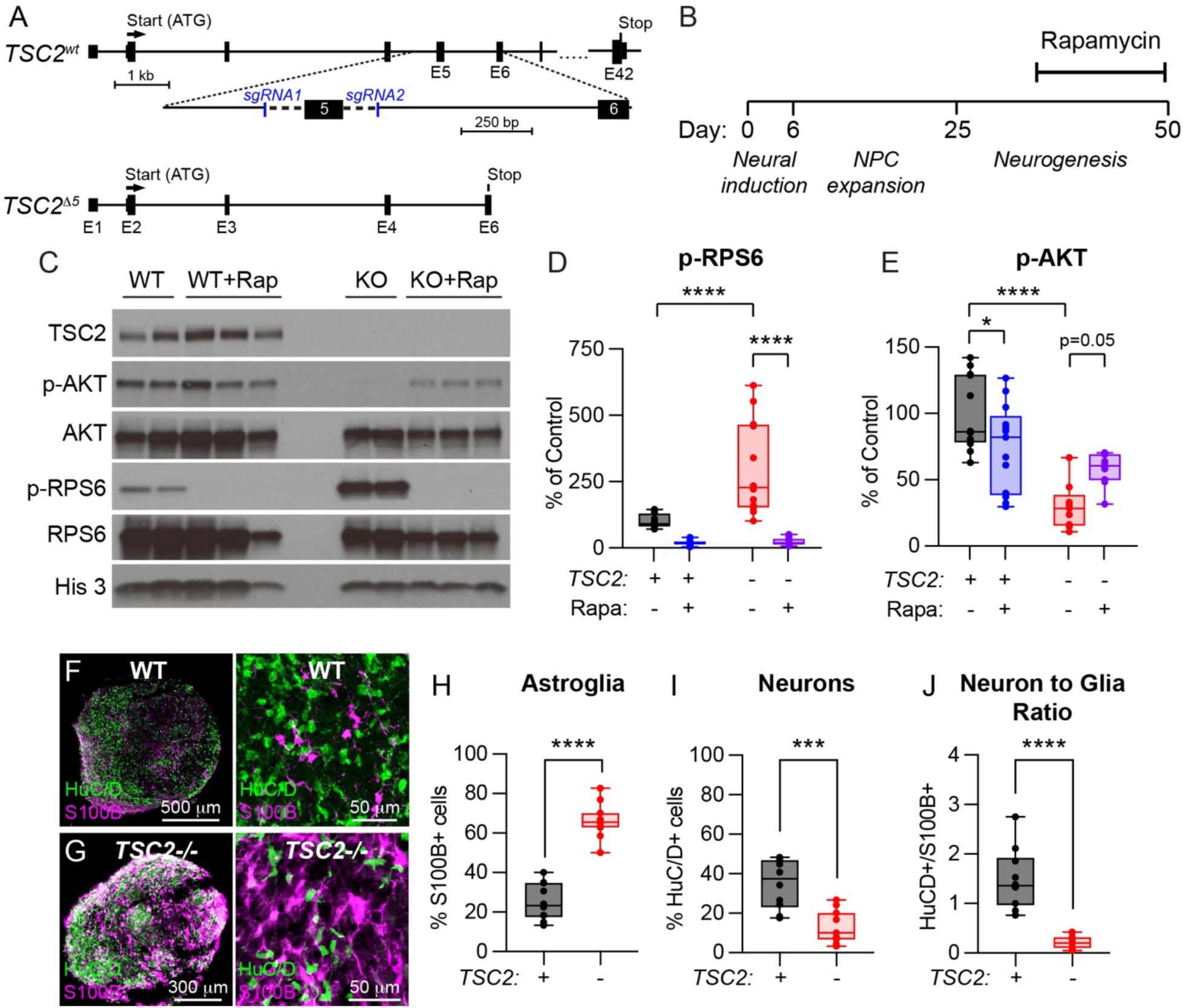
*TSC2^-/-^* human brain organoids have elevated mTORC1 signaling and reduced neuron to glia ratio. **(A)** Schematic of the CRISPR/Cas9-based editing approach to disrupt *TSC2* expression. Two guide RNAs were designed to target either of side of exon 5. Removal of exon 5 results in loss of TSC2 expression. **(B)** Timeline of the experiment. 50 nM rapamycin was added on day 35 of differentiation and organoids were harvested for western blot on day 50. **(C)** Example western blots of brain organoid lysates for the indicated conditions. 2-3 independent organoid samples per treatment are shown. **(D)** Western blot quantification for p-RPS6 (Ser240/244) normalized to total RPS6. N=12 organoids per condition. ****p<0.0001, one-way ANOVA; WT vs WT+Rap p=0.0962, WT vs KO ****p<0.0001, KO vs KO+Rap ****p<0.0001, Sidak’s multiple comparisons tests. **(E)** Western blot quantification for p-AKT (Ser473) normalized to total AKT. WT+vehicle n=12, WT+Rapa n=13, KO+vehicle n=9, KO+Rapa n=10 organoids. ****p<0.0001, one-way ANOVA; WT vs WT+Rap *p=0.0410, WT vs KO ****p<0.0001, KO vs KO+Rap p=0.0521, Sidak’s multiple comparisons tests. **(F)** Image of a section of a D100 WT brain organoid immunostained with HuC/D (green) and S100B (magenta). Right panel shows a zoomed-in image. **(G)** Image of a section of a D100 *TSC2* KO brain organoid immunostained with HuC/D (green) and S100B (magenta). Right panel shows a zoomed-in image. **(H)** Box and whisker plot of the % S100B+ nuclei for WT and *TSC2* KO organoids. ****p<0.0001; unpaired two-tailed t test. N=10 WT and 11 KO organoids **(I)** Box and whisker plot of the % HuC/D+ nuclei for WT and *TSC2* KO organoids. ***p=0.0001; unpaired two-tailed t test. N=10 WT and 11 KO organoids. **(J)** Box and whisker plot of the ratio of HuC/D+ to S100B+ nuclei for WT and *TSC2* KO organoids. ****p<0.0001; unpaired two-tailed t test. N=10 WT and 11 KO organoids. Boxes extend from the 25^th^ to 75^th^ percentiles, whiskers extend from the min to max values, lines represent the median, and dots represent values for individual organoids. For panels H-J, values are the same as for GFP-cells in shControl-treated organoids in Fig. 1H-J (WT) and Fig. 2C-E (KO).

**Supplementary Figure 2:**
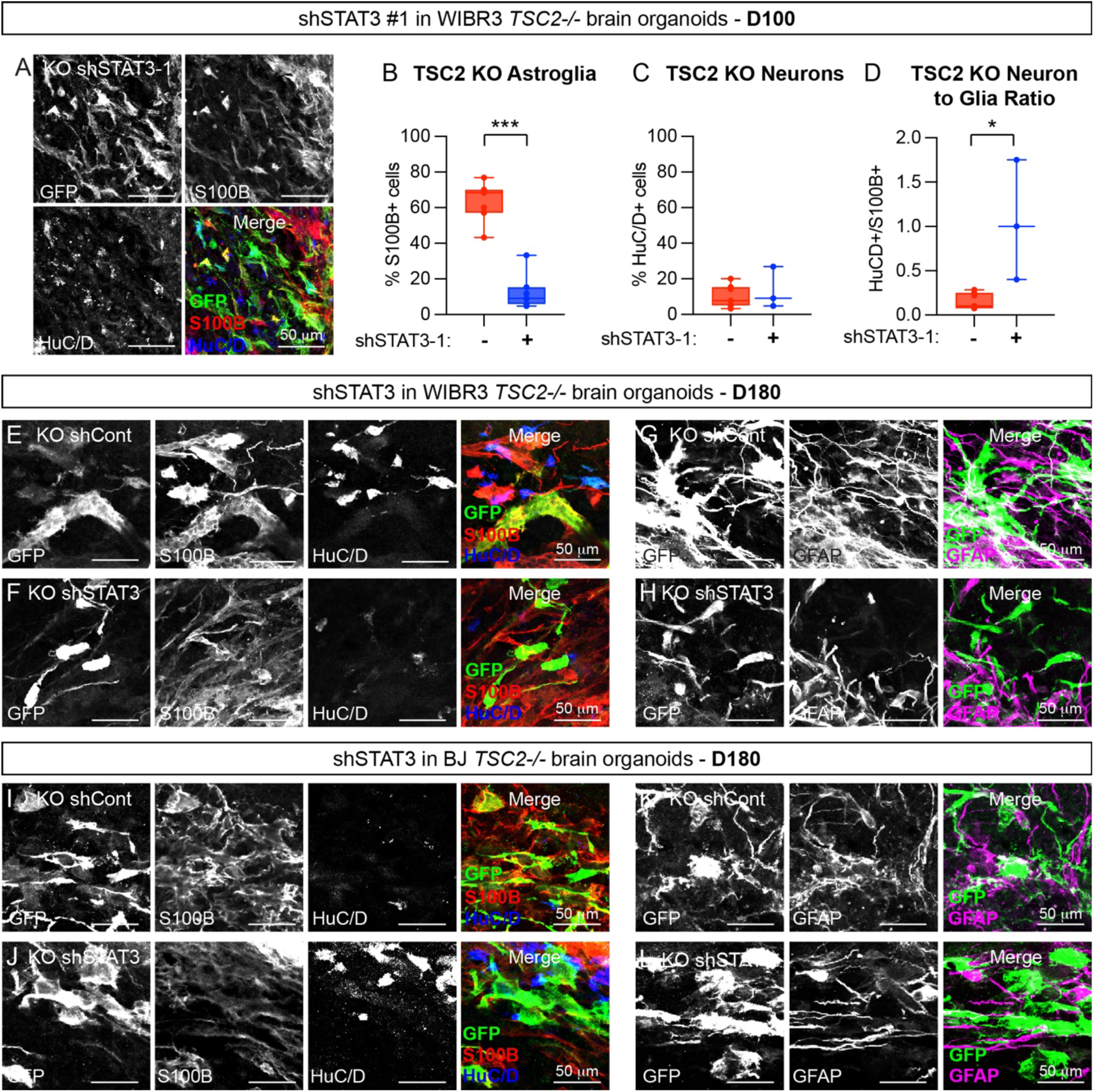
shSTAT3 reduces astrogenesis in *TSC2^-/-^* brain organoids. **(A)** Images of a day 100 WIBR3 *TSC2-/-* section from an organoid treated with shSTAT3-1-GFP and immunostained for S100B and HuC/D. **(B)** Box and whisker plot of the % S100B+ nuclei for GFP- and shSTAT3-1-GFP+ cells. ***p=0.0006, Mann-Whitney test, n=7 organoids. **(C)** Box and whisker plot of the % HuC/D+ nuclei for GFP- and shSTAT3-1-GFP+ cells. p=0.8333, Mann-Whitney test, n=7 organoids for GFP-cells and 3 organoids for GFP+ cells. **(D)** Box and whisker plot of the ratio of HuC/D+ to S100B+ cells. *p=0.0167, Mann-Whitney test, n=7 organoids for GFP-cells and 3 organoids for GFP+ cells. Boxes extend from the 25^th^ to 75^th^ percentiles, whiskers extend from the min to max values, lines represent the median, and dots represent values for individual organoids. For C and D some organoid sections had zero HuC/D+/shSTAT3-1-GFP+ cells and were therefore not included in the analysis. **(E,F)** Images from WIBR3 *TSC2-/-* organoids at D180 showing shControl-GFP labeled **(E)** or shSTAT3-GFP labeled **(F)** cells with S100B and HuC/D immunostaining. **(G,H)** Images from WIBR3 *TSC2-/-* organoids at D180 showing shControl-GFP labeled **(G)** or shSTAT3-GFP labeled **(H)** cells with GFAP immunostaining. **(I,J)** Images from BJ *TSC2-/-* organoids at D180 showing shControl-GFP labeled **(I)** or shSTAT3-GFP labeled **(J)** cells with S100B and HuC/D immunostaining. **(K,L)** Images from BJ *TSC2-/-* organoids at D180 showing shControl-GFP labeled **(K)** or shSTAT3-GFP labeled **(L)** cells with GFAP immunostaining.

**Supplementary Figure 3:**
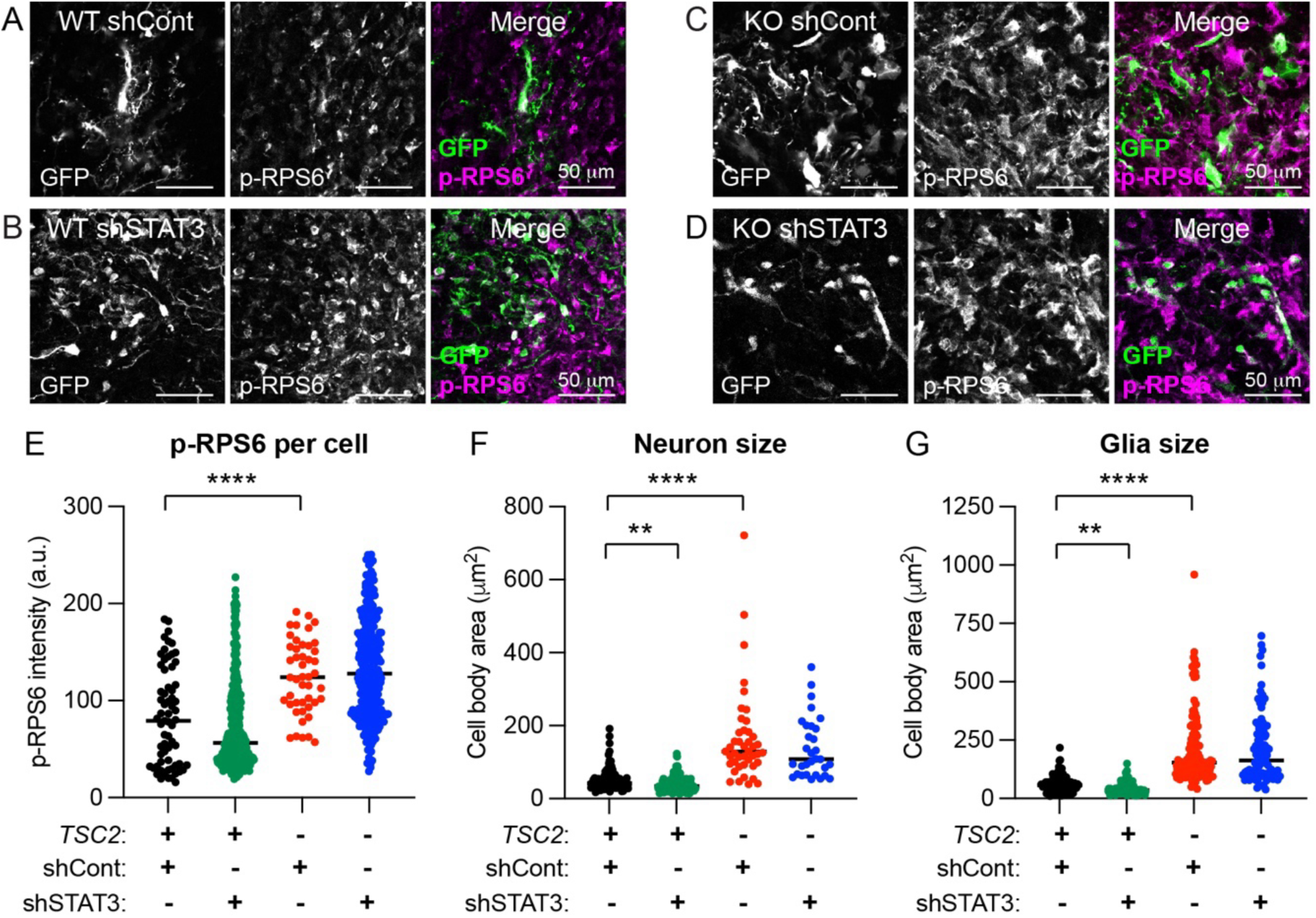
Reduction of STAT3 does not reverse mTORC1 hyperactivity in *TSC2^-/-^* cells. **(A-D)** Example images of sections from WT (**A,B**) and *TSC2-/-* (**C,D**) D100 brain organoids immunostained for p-RPS6 (Ser240/244). **A** and **C** show shControl-GFP transduced cells and **B** and **D** show shSTAT3-GFP transduced cells. **(E)** Scatterplot of p-RPS6 fluorescence intensity per cell for the indicated conditions. ****p<0.0001, Kruskal-Wallis test; WT/shControl vs WT/shSTAT3 p=0.4815, WT/shControl vs KO/shControl ****p<0.0001, KO/shControl vs KO/shSTAT3 p>0.9999, Dunn’s multiple comparisons test. **(F)** Scatterplot of neuron (HuC/D+) cell body area for the indicated conditions. ****p<0.0001, Kruskal-Wallis test; WT/shControl vs WT/shSTAT3 p=**0.0018, WT/shControl vs KO/shControl ****p<0.0001, KO/shControl vs KO/shSTAT3 p>0.9999, Dunn’s multiple comparisons test. **(G)** Scatterplot of astroglia (S100B+) cell body area for the indicated conditions. ****p<0.0001, Kruskal-Wallis test; WT/shControl vs WT/shSTAT3 p=**0.0061, WT/shControl vs KO/shControl ****p<0.0001, KO/shControl vs KO/shSTAT3 p>0.9999, Dunn’s multiple comparisons test.

**Supplementary Figure 4:**
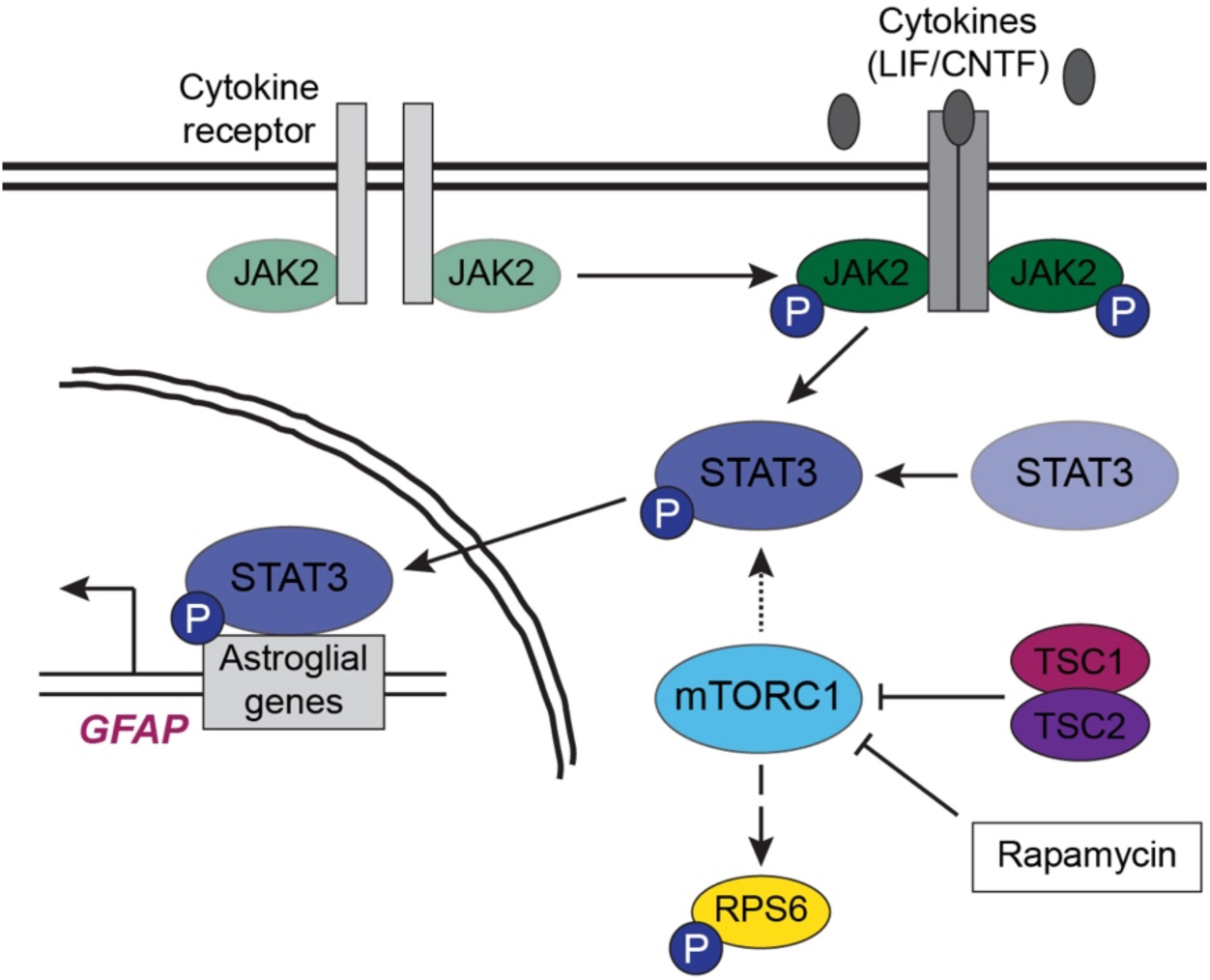
Summary of main findings. This study, together with prior work, shows that constitutively active mTORC1 signaling due to loss of negative regulation by the TSC1/2 complex leads to enhanced STAT3 phosphorylation. This, in turn, drives increased astrogenesis in human brain organoids. By contrast, mTOR inhibition with rapamycin reduces astrogenesis and concomitantly increases neurogenesis. Treatment with the cytokines LIF and CNTF induces STAT3 phosphorylation and GFAP expression. These effects do not require mTORC1 signaling as they are not blocked by rapamycin treatment. The mechanism by which mTORC1 regulates STAT3 phosphorylation during brain development is currently unknown (indicated by the dashed arrow).

**Supplementary Table 1:** Array Comparative Genomic Hybridization (aCGH) analysis of hESC and hiPSC lines.

| **Cell line** | **Chrom.** | **Amp/**<br>**Del** | **Start** | **Stop** | **Size (kb)** | **Chrom.**<br>**band** | **Log2**<br>**Ratio** |
| --- | --- | --- | --- | --- | --- | --- | --- |
| WIBR3<br>WT<br>hESCs | 20 | Amp | 31,300,674 | 32,533,660 | 1,232.987 | q11.21 | 0.900 |
| WIBR3<br>*TSC2*<sup>-/-</sup><br>hESCs | 20 | Amp | 31,300,674 | 32,533,660 | 1,232.987 | q11.21 | 0.508 |
| BJ<br>*TSC2*<sup>-/-</sup><br>hiPSCs | 7 | Amp | 5,351,046 | 7,097,949 | 1,746.904 | p22.1 | 0.537 |
|  | 15 | Del | 20,276,449 | 22,221,303 | 1,944.855 | q11.1 -<br>q11.2 | -1.140 |

## Notes

### Competing Interest Statement

The authors have declared no competing interest.

